# Lossless compression of k-mer matrices enabling random row access

**DOI:** 10.64898/2026.07.03.736306

**Authors:** Alix Regnier, Téo Lemane, Sébastien Bellenous, Rayan Chikhi, Pierre Peterlongo

## Abstract

Genomic search engines such as `Logan-Search` index petabytes of sequencing data as large binary matrices, called *k*-mer matrices, where each row encodes the presence of a *k*-mer across thousands to millions of genomic samples. `Logan-Search` contains a petabyte of binary matrices, and storing them is expensive, yet compression must not prevent fast random access to any matrix row at query time.

We present `kmcomp`, a lossless compression method for *k*-mer matrices that satisfies these competing requirements. Block compression partitions the matrix into fixed-size row blocks, each compressed independently; block start positions are stored in an `Elias-Fano` encoded array, enabling *O*(1) random access to any block. To improve compressibility without introducing additional decompression steps, we introduce the *π*-compression: a column reordering that groups similar samples together by solving the Traveling Salesman Problem via a nearest-neighbor heuristic. We accelerate this heuristic with a novel variant of the vantage-point tree, the `masked vp-tree`, which dynamically prunes nearest-neighbor search space.

On three (meta)genomic datasets, `kmcomp` achieves compression ratios of 1.3 to 5.4; *π*-compression further improves these to 1.5 to 51.3. Applied to the `Logan-Search` petabyte-scale index, compression reduces storage by approximately half, and *π*-compression adds a further 13% gain. Query overhead remains modest: queries of hundreds of nucleotides incur an absolute latency increase of *≈* 100 ms, and highly compressed indexes can match uncompressed query times thanks to reduced disk reads.

## 1 Introduction

Searching a genomic sequence against a petabyte-scale database, such as SRA [17], can nowadays be performed using genomic search engines such as `MetaGraph` [16] or `Logan-Search` [7]. Genomic search engines offer invaluable possibilities for researchers, as they enable researchers to estimate the distribution of a sequence of interest across millions of experiments, ultimately allowing easy correlations with metadata. Making petabyte-scale databases searchable is a new era in bioinformatics, as it brings unmatched knowledge for any homologous study.

Search engines rely on indexes, which are data structures that preprocess data to answer queries without parsing the entire database. In bioinformatics, such indexes usually rely on *k*-mers, which are overlapping substrings of size *k*. A query consists of retrieving genomic document IDs associated with a given *k*-mer. Storing such associations is an extensively studied field in bioinformatics. For interested readers, Marchet [20] gives an overview of the state of the art *k*-mer indexes.

`Logan` [7] project indexed more than 5 *×* 10^16^ basepairs across 27.3 million SRA accessions. As a result, `Logan-Search` relies on an index of 1 petabyte of *k*-mer matrices built with `kmindex` [19]. Given a set *S*= {*S*_1_, …, *S*_*N*_} composed of *N* samples, we call a “*k*-mer matrix” a row-major binary matrix *M* ∈ {0, 1} ^*N×m*^. In *M*, each row *i* ∈ [0, *N* [ represents the address of a given *k*-mer, and each column *j* ∈ [0, *m*[ represents *k*-mers from the sample *S*_*j*_. In practice, *m* can reach values in the billions.

The bit *M* [*i, j*] tells if the *k*-mer corresponding to the row *i* belongs to *S*_*j*_ or not. Hence, a row is a vector of booleans (also called a bitvector) that represents the colored set for its corresponding *k*-mer. Querying such a data structure consists of streaming *k*-mers from a given sequence, hashing them, and retrieving their corresponding bitvectors. This implies that both latency and throughput of row retrievals are crucial for keeping queries fast. Note that in the `Logan-Search` context, the columns are Bloom filters [4] using a unique hash function. This specificity does not affect the proposed work.

It combines the reuse of general-purpose compressors with novel, theoretical, and practical propositions. Precisely, we address the following problem.

### Problem 1

*Given an uncompressed binary matrix M* ∈ {0, 1} ^*N×m*^, *propose a lossless compressed version M*_*c*_ *of M, such that:*

1. *The compression method must be sufficiently fast to scale to petabytes of data. Compression must be orders of magnitude faster than index creation*.
2. *Random access to any row M*_*c*_[*i*] *of the compressed matrix does not require the decompression of the whole matrix*.
3. *Row retrievals on M*_*c*_ *are done in time of the same order of magnitude as on M* .

In summary, we make the following contributions:

– A block compression scheme for *k*-mer matrices with *O*(1) random row access, using `Elias-Fano` encoded block locations.
– *π*-compression: a column reordering heuristic that models the optimal ordering as a path TSP, and solves it via a novel `masked vp-tree`, achieving *O*(log *N* ) average nearest-neighbor queries without replacement.
– The *π*_gain_ metric, computable in *O*(*N* ) distance evaluations before any reordering, to predict whether *π*-compression will be beneficial for a given matrix.
– Experiments on three genomic collections and the petabyte-scale `Logan-Search` index, showing compression ratios up to 51*×* and halving the storage of `Logan-Search`.

Note that our contribution is initially motivated by questions related to *k*-mer indexing, but it is not specific to this purpose. The concepts proposed here apply to compressing any binary matrix whose columns exhibit similarity.

## 2 Related works

Several compressed representations of large binary matrices have been proposed, particularly in graph and annotation storage. The depth-first representation of *k*^2^-trees [6] compresses sparse binary relations through recursive matrix decomposition. Although highly space-efficient, row retrieval requires traversing multiple levels of the tree structure, which introduces prohibitive latency.

The `RowDiff` technique [8] exploits the fact that in a colored de Bruijn graph, consecutive *k*-mers tend to share similar colored sets. However, `Logan-Search` *k*-mer matrices use a random hash function, which makes consecutive bitvectors independent. Thus, `RowDiff` is inappropriate for compressing such *k*-mer matrices.

Rahman et al. [21] proposed a compression algorithm for colored de Bruijn graph *k*-mer colored sets. While yielding good compression results, it does not provide random *k*-mer colored set access without complete decompression, making such a technique unsuitable for answering Problem 1.

Method `OnPair` [12] addresses a related problem. It is designed to compress a sequence of short strings while enabling fast random access, and *k*-mer matrix rows could be considered as strings. However, `OnPair` relies on Longest Prefix Matching processes. This is computationally too expensive when strings are not similar. In this work, rows are independent, which precludes the use of such a method.

To the best of our knowledge, this work fills a gap, as there exists no method dedicated specifically to compressing large binary matrices while offering random access to any row in the compressed matrix in parametrized constant time.

## 3 Methods

Figure 1 outlines the `kmcomp` pipeline. The key contributions lie in the first two panels and the block compression algorithm detailed in the last panel, each described in the sections below.

**Fig. 1:**
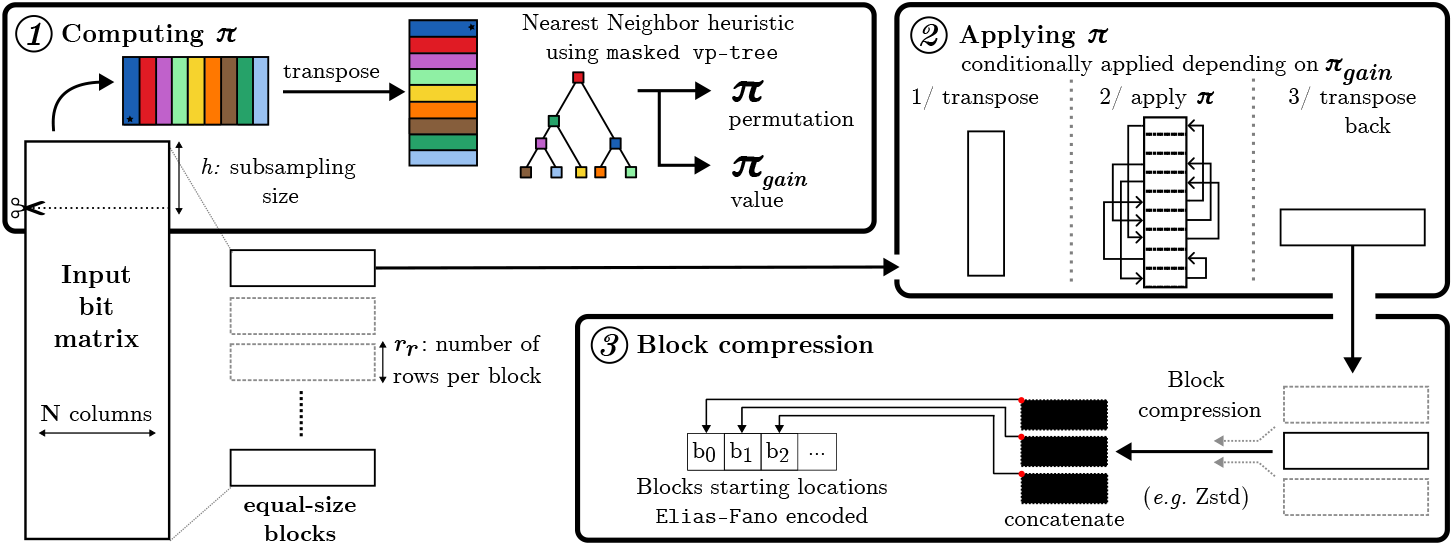
Methods overview. *①* Computation of the permutation *π* and of the metric *π*_gain_ by combining subsampling of matrix rows and `masked vp-tree` on transposed columns. *②* Application of permutation *π* with double block transposition, this stage is conditionally applied if a given threshold on *π*_gain_ is satisfied. The bit matrix is split into blocks; each block is transposed, its rows (transposed columns) are reordered according to *π*, and it is transposed back. *③* Block compression is immediately executed after stage *②* (or after stage *①* if stage *②* is not applied), any block is compressed on the fly, and its new starting location is stored in an `Elias-Fano` encoded vector.

### 3.1 Block compression

The main issue with row compression is that the exact size of any compressed row cannot be determined without decompressing it or explicitly recording it. One might think that we could simply compress each row, concatenate the compressed representations, and store their positions sequentially. Although this idea provides random access to any compressed row, it has major drawbacks. First, *k*-mer matrices can have billions of rows. Storing row offsets thus incurs substantial memory overhead. Second, a single row may not be compressible, as it provides little context for compression. As a solution, we propose compressing rows in blocks. Block compression is shown in panel *③* of Figure 1.

While widely adopted by many general-purpose compressors such as XZ^4^ or Zstandard^5^, block compression consists of breaking data into blocks of the same size. These blocks can be compressed (and decompressed) in parallel. The block size is a trade-off between time and output size. Smaller size reduces the time required to process a block, but it also reduces the context size, leading to lower compression (as depicted in Figure S1 in supplementary materials).

#### Block random access

We borrow the “block compression” concept from general-purpose compressors to compress *k*-mer matrices into blocks containing an arbitrary number of rows. Each block is compressed independently, preventing the introduction of dependency constraints during decompression. Compressed blocks are concatenated, and the starting position of each block is stored in an array *L*. In order to delimit the last block, the cumulative size of compressed blocks |*M*_*c*_| is pushed to the end of *L*. With *n* being the number of compressed blocks and *b*_*i*_ the starting location of the *i*-th block, we have *L* = [*b*_0_, *b*_1_, …, *b*_*n* 1_, |*M*_*c*_| ]. For any block to be decompressed, *L* provides its start and end locations. This information is used for decompressing any block. As *L* is a sequence of strictly increasing integers, it is encoded using `Elias-Fano` [11, 9]. This succinct data structure provides random access to any block location in **O**(1) time.

#### Retrieving a row

Retrieving a row whose index is *x* requires decompressing the block 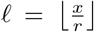, and then accessing the row at address *x* mod (*r*) in the decompressed block. Note that retrieving a single row involves decompressing the full block to which the row belongs. This slows down queries. Sorting the queried row indices in advance groups all queries within a block into a single sequential scan, avoiding repeated decompression of the same block.

Therefore, our strategy is to keep the last decompressed block in memory, since the next row to be queried is likely to belong to the same block. Hence, each block is decompressed at most once at query time.

As a side note, one might think that to improve compression, we just need to increase the block size. But in reality, such an idea usually yields poor compression gains (theoretically on the order of a logarithm) at the cost of additional compression time (theoretically linear).

### 3.2 Column reordering before compression

The compression efficiency depends heavily on whether similar items are colocated before compression. This idea notably inspired the so-called “phylogenetic compression” [5, 14], which uses evolutionary history to reorder a set of genomes to be compressed. In the binary matrices we manipulate, the original column order is arbitrary. It can be modified to group similar columns together, reducing data complexity by introducing longer runs of bits. Most window-based compression techniques would benefit from this reordering, since redundancies could be captured together within a small window.

As a way to maximize the number of bits in runs of 0s and in runs of 1s in the rows of the matrix, we suggest a heuristic for grouping columns of the matrix by similarity. The objective is to 1/ determine a permutation *π* of ⟦0; *N −*1⟧ of the columns and 2/ apply this permutation efficiently. Determining *π* can be modeled using a complete graph in which vertices are matrix columns, connected by weighted edges representing “*similarity*” of each pair of matrix columns. The optimal *π* permutation is then obtained through a Traveling Salesman Problem (TSP) of this graph, equivalent to finding a Hamiltonian path minimizing the sum of edge weights. The problem is known to be NP-hard [13]. Several techniques can serve as heuristics for computing *π*, such as Neighbor-Joining [22], which is used in phylogenetic compression. However, none of these techniques can beat the simplicity and efficiency of the Nearest Neighbor heuristic. As illustrated Figure 1, panel *②*, we propose an algorithm that achieves an efficient approximation of *π* by coupling Nearest Neighbor with a metric tree [23] such as the vantage-point tree (`vp-tree`) [24]. Finally, in Section S2 we give technical details of how efficiently applying *π* to a row-major matrix and why it matters.

### 3.3 On faster Nearest Neighbor heuristic

To the best of our knowledge, TSP computation algorithms assume that the distance between any pair of elements is given as input or is computed in *O*(1) time complexity. Within the present framework, this does not hold because columns can extend to several billion bits, and computing the distance between two columns is time-consuming. In practice, column proximity is measured by the Hamming distance. In Section S3, we provide further details on why we use this distance and how it is efficiently computed.

To reduce distance computation time, we propose two solutions. First, we compute the distance of columns using column subsampling (see Section S4). Second, we prune the search space for nearest-neighbor queries. However, given the large number of dimensions, we cannot rely on multidimensional search trees such as `kd-tree` [2]. As a solution, we use a vantage-point tree.

#### The vantage-point tree

The vantage-point tree (`vp-tree`) [24] is both a binary and metric tree [23]. This data structure aims to find the nearest neighbor of an element *l* with a branch-and-bound algorithm that converges by updating the current vertex *α* whose distance to *l* is minimal among all elements except *l*. We give vp-tree time complexities in Section S5.

#### The masked vantage-point tree

Since the TSP path formulation forbids vertex revisits, we introduce a modification to `vp-tree`, named “`masked vp-tree`”. Let *π* be a queue, initialized with a randomly chosen column. Iteratively, we push the nearest neighbor of the last element of *π* to *π*. To prevent `vp-tree` from converging to a vertex that was already added (already visited) to *π*, we do not update *α* when we encounter a vertex that is already visited (see Algorithm S1). This modification enables the algorithm to converge toward the closest unvisited neighbor and gradually prune the search space. To this end, we added a “*masked*” flag to each `vp-tree` vertex.

##### Definition 1

(Masked node in a `vp-tree`)

*A node in the* `vp-tree` *is “masked” if and only if the two following conditions are satisfied:*

– *its associated vertex is on the path in the TSP*.
– *has no children or all children are masked*.

To implement Definition 1, we need to check if a vertex is masked. It implies checking if the vertex has been added to *π* and if all subtree vertices are masked.

Checking whether a vertex has been added to *π* is done in constant time via a bitvector lookup, but checking each subtree vertex would require *O*(*N* ) lookups, which would be inefficient. By storing a masked flag for each node, we can know if all subtree vertices are masked in constant time by checking if the flag for each of its direct children is set (*i*.*e*. two checks). Such flags are updated each time a vertex is added to *π* by recursively updating masked flags from the added vertex to the root (bottom-up). This update procedure (see Algorithm S2) is stopped as soon as the current vertex cannot be masked, as neither can be its following ancestors (as established in Definition 1). This procedure has *O* (*N* ) space complexity, and updates take *O* (log *N* ) time. Overall, `masked vp-tree` time and space complexities are the same as a `vp-tree`.

### 3.4 Predicting when it is worth reordering columns

Reordering the columns is a costly task. Therefore, one needs a way to predict the expected compression gain from reordering. The objective is to estimate the number of bit-runs, in the original matrix compared to those found if the matrix were reordered. A proxy of this metric is to sum the probabilities of breaking a bit-run among all consecutive columns, which gives the number of expected bit-runs in a row. In other words, we sum the Hamming distances of all consecutive columns in the original matrix and as it would be in the reordered matrix. For any matrix, we note *c*[*j*] as the *j*-th column of the matrix and *c*[*π*[*j*]] as the new *j*-th column of the matrix if a given permutation *π* was applied. We divide these two sums to compute a metric that we call “*π*_gain_” (see Equation (1), with *d*_*H*_ standing for the Hamming distance). This metric estimates how much reordering might improve the lengths of bit-runs after reordering. It is important to notice that when *π* is known, one can compute *π*_gain_ in *O*(*N* ) distance computations, before *π* is applied to the matrix.

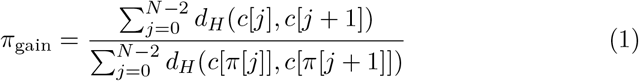

## 4 Results

We exploit the practical advantages of `kmindex` namely its compact indexes, straightforward usage, and scalability to massive datasets. The following experiments are therefore intentionally scoped to demonstrate `kmcomp` on `kmindex` outputs, rather than to compare indexing tools.

### 4.1 Compressor choice

We selected Zstandard^6^ (Zstd) because it provides one of the best compromises between compression ratio, compression speed, and decompression speed among general-purpose compressors. Moreover, Zstd is very efficient for decompressing data with low latency. Such properties are crucial for a fast decompression of independent blocks, as required by the block compression scheme.

### 4.2 Datasets

#### Experimental datasets

We used the same datasets as in Fan et al. [10] to study the effect of the parameters of `kmcomp`. Metrics of datasets are provided in Table 1. They were indexed using `kmindex` [19].

**Table 1:**
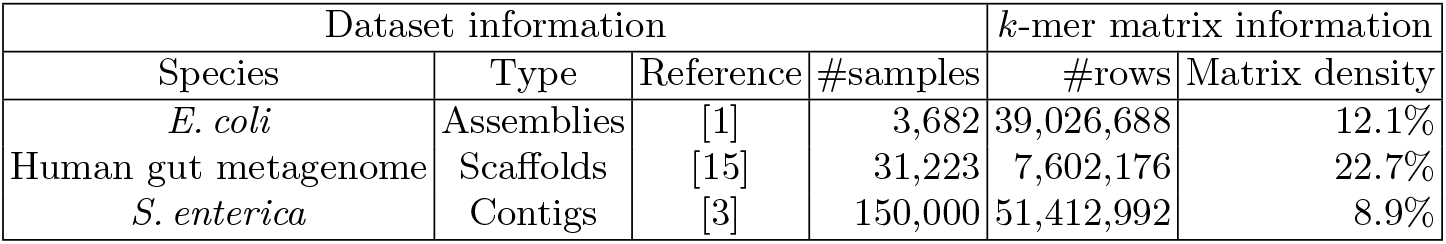
Experimental datasets information. Matrix density provides the ratio of 1’s over the total matrix size in matrices created by kmindex on these datasets.

#### Logan-Search index *k*-mer matrices

Petabyte-scale experiments were done on a subpart `Logan-Search` [7]. `Logan-Search` is an indexation of a Sequence Read Archive (SRA) snapshot, totaling about 27 million SRA accessions and about 5 *×* 10^16^ basepairs. `Logan-Search` is built over 1 petabyte of *k*-mer matrices, which were also created using the `kmindex` tool. SRA accessions were split into 2869 sub-indexes based on multiple factors: (i) the omic level (genomic, transcriptomic, metagenomic, … ) ; (ii) the superkingdom, which was automatically attributed using STAT [18] (see Table S2) ; (iii) the log of the estimated number of distinct *k*-mers. In the following experiments, *π* has been computed once for each sub-index.

### 4.3 Main results

The implementation repositories and the experiment scripts repository are described in Sections S6 and S7, also providing hardware specifications.

Reported query times are measured via `kmindex`, which pre-sorts queried *k*-mers to avoid cache misses and limit decompression to at most one pass per compressed block.

#### Compression size on experimental datasets

Table 2, shows compression sizes for *E. coli*, Human gut metagenome, and *S. enterica* datasets. Compression ratios of about 4.1, 1.3, and 5.4, respectively. When indexes are *π*-compressed, compression improves by up to *≈*10 times, yielding compression ratios of 9.3, 1.5, and 51.3, respectively. The compression on the Human gut metagenome dataset is less effective than on the other datasets. This is expected, as this dataset is composed mostly of low-related species and is therefore less compressible. Our hypothesis is that such datasets are less prone to *π*-compression.

**Table 2:** Index size whether matrices are not compressed, compressed, or *π*-compressed and query time (average and standard deviation for 5 replicas) whether matrices are not compressed or *π*-compressed. Block size was set to 64 KB. Subsampling size was set to 10,000.

|  | Dataset<br>Method | *E. coli* | Human gut<br>metagenome | *S. enterica* |
| --- | --- | --- | --- | --- |
| Index size (GB) | no comp. | 16.8 | 27.6 | 897.8 |
|  | compressed | 4.1 | 21.1 | 167.2 |
| | π-compressed | 1.8 | 17.7 | 17.2 |
| Query time (s) **mean ± std**<br>query size=150 nt | no comp. | 0.02 ±0.00 | 0.06 ±0.00 | 0.34 ±0.02 |
| | π-compressed | 0.13 ±0.00 | 0.17 ±0.00 | 0.34 ±0.02 |
| Query time (s) **mean ± std**<br>query size=150,000 nt | no comp. | 1.78 ±0.09 | 8.14 ±0.12 | 36.00 ±0.74 |
| | π-compressed | 9.22 ±0.10 | 28.00 ±0.10 | 40.00 ±0.73 |

In Section S8, we provide further details on *π*_gain_ values for distinct datasets. We give details on how compression behaves when varying the size of blocks in Section S1.

#### Compression efficiency

We note as “compression efficiency”, the amount of data that a method can save per second during the compression steps (see formulas in Section S9). Compression efficiency results are shown Table 3.

**Table 3:** Saved bits per second at compression time, whether indexes are compressed or *π*-compressed. Subsampling size was set to 10,000. Block size was set to 64 KB. Repeated 8 times.

| Dataset | Saved MB/s **mean ± std** |  |
| --- | --- | --- |
| | compressed | π-compressed |
| *E. coli* | 144.4 ±03.4 | 151.1 ±14.3 |
| Human gut metagenome | 76.4 ±52.0 | 27.0 ±02.9 |
| *S. enterica* | 298.3 ±07.7 | 382.8 ±09.2 |

In the case of the Human gut metagenome dataset, *π*-compression is *>* 4 times longer than compressing without reordering and yields a compression improvement of about 19%. Overall, in this case, reordering results in a loss of compression efficiency. For the genomic datasets, *π*-compression improves the efficiency when compared to compression efficiency without reordering.

In Section S10, we give further details on the time spent for each stage of the pipeline.

#### Time for computing *π* on experimental datasets

Figure S5 shows that `masked vp-tree` outperforms `vp-tree` for solving the path TSP Nearest-Neighbor heuristic in all tested cases. However, `vp-tree` approaches can be less efficient than a naive approach. On the Human gut metagenome, `vp-tree` is 40% slower than a naive approach. In worst-case scenarios, `vp-tree` and masked `vp-tree` seem to end up computing more pairwise distances than the naive approach (see Figure S6). On the *E. coli* and *S. enterica* datasets, `masked vp-tree` is faster than the naive approach by factors of 8 and 170, respectively. When comparing Figure S5 and Figure S6, we remark that the *π* computation time for `vp-tree` and `masked vp-tree` correlates with the number of computed distances.

These experiments were conducted with default parameters. We present additional results in Section S11 as we vary the subsampling size.

#### Query time on experimental datasets

As shown Table 2, for queries composed of 150 nt, compressed indexes answer queries with a time penalty of about 100 ms. For queries of 150,000 nt, we report that query times slowed down by a factor of 1.1 to 5.2 on compressed indexes. However, on the *S. enterica* dataset, we observe that query time is the same regardless of whether the index is compressed. *S. enterica* index is about 50 times smaller when *π*-compressed, which reduces disk I/O, resulting in faster queries. We additionally present results when varying the block sizes in Section S12.

#### Compression of Logan-Search index

The objective is to estimate the expected compression of the complete index. For each `Logan-Search` sub-index, we applied our pipeline to a single matrix (each index is subdivided into 256 matrices composed of the original *N* columns and 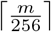 rows). We extrapolate such results to estimate how compressible `Logan-Search` index *k*-mer matrices are and how much compression can be done via *π*-compression. We computed *π* for each `Logan-Search` sub-index, in 2 days of CPU time. We estimate that complete *π*-compression would take 125 days of CPU time. Note that the full construction of `Logan-Search` took 467,200 days of CPU time. Hence the compression itself would be only 0.3% of the total index creation.

As shown in Table 4, compression alone, without column permutations, would reduce the index size from approximately one petabyte to approximately 500 terabytes. Using *π*-compressed would improve compression by about 13%. Further interpretation is provided in the Discussion Section.

**Table 4:** Logan-search estimated index size whether the index is unchanged, compressed, or *π*-compressed.

| Logan-search estimated index size (TB) |  |  |
| --- | --- | --- |
| no compression | compressed | π-compressed |
| 1023.8 | 514.2 | 455.3 |

#### Predicting the effect of column reordering on compression

Figure S11 shows that *π*_gain_ strongly correlates with the actual compression gain yielded by reordering (*R*^2^ *≈* 0.94). This makes *π*_gain_ a reliable predictor of the benefit of applying *π*.

## 5 Discussion

We proposed `kmcomp`, a compression tool for large binary matrices that enables fast random access to any row in the compressed representation. With the so-called *π*-compression, `kmcomp` takes advantage of potential column similarities by reorganizing columns in similarity order before compression. Because column reordering is computationally expensive, we introduce a metric to estimate the potential compression gain before performing the reordering. Experimental results show that this metric is data-invariant and provides accurate estimates.

Computing the column permutation entails solving an approximate TSP over thousands to millions of columns, minimizing the total distance between each pair of consecutive columns. In this context, we introduced the `masked vp-tree` data structure, which enables faster nearest-neighbor search and on which the TSP nearest-neighbor heuristic is based. We show that `masked vp-tree` can be used to efficiently pick nearest neighbors without replacement with a logarithmic update complexity. Such a data structure can be used to find the exact nearest neighbor of a given element when the number of dimensions is too high to use data structures subject to the curse of dimensionality, like `kd-tree` [2]. It is worth noting that it might be interesting to see whether using approximate nearest neighbors by relaxing the `masked vp-tree` branch-and-bound would speed up the construction of *π* without compromising compression.

We developed `kmcomp` in the context of bioinformatics to compress *k*-mer matrices, where each row represents the presence/absence of *k*-mers in a genomic dataset. Querying a nucleic sequence of size *l* implies accessing *l − k* + 1 rows. `kmcomp` was originally developed to compress the `Logan-Search` index, a collection of such matrices totaling 1 PB.

We have shown that *π*-compression improves *k*-mer matrix compression by factors of 1.19 to 9.48 compared to the original compression. We have systematically found an equal or better order for the tested *k*-mer matrices by solving it as an approximate TSP. Furthermore, `kmcomp` could be extended beyond bioinformatics to any binary matrices indexing related documents.

Querying a compressed matrix can be as fast as querying an uncompressed one, but in some cases it can take up to 6 times longer. This is notably the case of poorly compressible datasets, in which columns exhibit little similarity, as when original data are divergent metagenomic samples. However, for queries of about a hundred nucleotides, we report an absolute slowdown of about 100 milliseconds compared to querying uncompressed matrices. In practice, the overhead when querying sequences of a few thousand nucleotides, as is the case in `Logan-Search`, is a few seconds.

Future work concerns the compressor itself, currently being Zstandard. As a byte-level compression algorithm, it does not fully exploit runs of 0s or 1s within each row. We will propose a Bit-level Run-Length encoder (BRLE) to compress bit-runs, which will also make *π*_gain_ even more accurate because such a compression scheme only depends on the length of bit-runs. Moreover, because BRLE is context-free, it could use smaller compressed blocks than the current framework does. Smaller blocks would, in turn, require less data to decompress when accessing a row, thereby improving query speed.

Regarding `Logan-Search` *k*-mer matrices, it is estimated that the cumulative size of the search engine can be reduced by more than half by applying `kmcomp`. We report that *π*-compression on such *k*-mer matrices further improved compression by 13%. We have identified several possible reasons why *π*-compression is less efficient than expected on this dataset. First, genomic documents were clustered into sub-indexes using a classification algorithm [18]. As shown Figure S12, many classifications were incorrect, and several unrelated species were misclassified into the wrong taxonomic division, limiting column similarities. Second, genomic sequences uploaded to the SRA are not error-free and were sequenced using several different technologies, suggesting that two genomic documents intended to index two closely related species can differ substantially. Further improvements in compression for `Logan-Search` could be achieved by clustering genomic documents more effectively.

Our current framework does not support appending new samples to a compressed *k*-mer matrix because `Zstd` does not support updates. Thus, future improvements to the compression algorithm will be designed to accommodate row extensions in the matrices.

In this work, we assume the hash function used to construct the *k*-mer matrix is predetermined. However, when constructing the *k*-mer matrix, it would be beneficial to use a Locality-Sensitive Hashing (LSH) function such as IDL [25]. Doing so would implicitly group consecutive *k*-mers that are more likely to share the same set of genomic documents, thereby clustering similar rows together and improving overall compressibility.

## Acknowledgments

This work was supported by state funding managed by the French National Research Agency under the France 2030 program [ANR-22-PEAE-0005], and by Inria challenge program OmicFinder. We acknowledge the GenOuest bioinformatics core facility for providing the computing infrastructure.

The authors acknowledge the Texas Advanced Computing Center (TACC) for providing computational resources that have contributed to the research results reported within this paper.

## Footnotes

4 https://github.com/tukaani-project/xz

5 https://github.com/facebook/zstd

6 https://github.com/facebook/zstd

## Bibliography

[1] Alanko, J.N.: 3682 e. coli assemblies from ncbi (2022), 10.5281/zenodo.6577997, URL https://doi.org/10.5281/zenodo.6577997

[2] Bentley, J.L.: Multidimensional binary search trees used for associative searching. Commun. ACM 18(9), 509517 (Sep 1975), ISSN 0001-0782, 10.1145/361002.361007, URL https://doi.org/10.1145/361002.361007

[3] Blackwell, G.A., Hunt, M., Malone, K.M., Lima, L., Horesh, G., Alako, B.T., Thomson, N.R., Iqbal, Z.: Exploring bacterial diversity via a curated and searchable snapshot of archived dna sequences. PLoS biology 19(11), e3001421 (2021), URL http://ftp.ebi.ac.uk/pub/databases/ENA2018-bacteria-661k

[4] Bloom, B.H.: Space/time trade-offs in hash coding with allowable errors. Communications of the ACM 13(7), 422–426 (1970)

[5] Břinda, K., Lima, L., Pignotti, S., Quinones-Olvera, N., Salikhov, K., Chikhi, R., Kucherov, G., Iqbal, Z., Baym, M.: Efficient and robust search of microbial genomes via phylogenetic compression. Nature Methods 22(4), 692–697 (2025)

[6] Brisaboa, N.R., Ladra, S., Navarro, G.: k2-trees for compact web graph representation. In: International symposium on string processing and information retrieval, pp. 18–30, Springer (2009)

[7] Chikhi, R., Lemane, T., Loll-Krippleber, R., Montoliu-Nerin, M., Raffestin, B., Camargo, A.P., Miller, C.J., Fiamenghi, M.B., Agustinho, D.P., Majidian, S., et al.: Logan: planetary-scale genome assembly surveys lifes diversity. bioRxiv pp. 2024–07 (2025)

[8] Danciu, D., Karasikov, M., Mustafa, H., Kahles, A., Rätsch, G.: Topology-based sparsification of graph annotations. Bioinformatics 37(Supplement_1), i169–i176 (2021)

[9] Elias, P.: Efficient storage and retrieval by content and address of static files. Journal of the ACM (JACM) 21(2), 246–260 (1974)

[10] Fan, J., Khan, J., Singh, N.P., Pibiri, G.E., Patro, R.: Fulgor: a fast and compact k-mer index for large-scale matching and color queries. Algorithms for Molecular Biology 19(1), 3 (2024)

[11] Fano, R.M.: On the number of bits required to implement an associative memory. Massachusetts Institute of Technology, Project MAC (1971)

[12] Gargiulo, F., Venturini, R.: Onpair: Short strings compression for fast random access. arXiv preprint arXiv:2508.02280 (2025)

[13] Hartmanis, J.: Computers and intractability: a guide to the theory of np-completeness (michael r. garey and david s. johnson). Siam Review 24(1), 90 (1982)

[14] Hendrychová, V., Břinda, K.: Why phylogenies compress so well: combinatorial guarantees under the infinite sites model. bioRxiv pp. 2026–03 (2026)

[15] Hiseni, P., Rudi, K., Wilson, R.C., Hegge, F.T., Snipen, L.: Humgut: a comprehensive human gut prokaryotic genomes collection filtered by metagenome data. Microbiome 9(1), 165 (2021), URL https://arken.nmbu.no/~larssn/humgut/

[16] Karasikov, M., Mustafa, H., Danciu, D., Kulkov, O., Zimmermann, M., Barber, C., Rätsch, G., Kahles, A.: Efficient and accurate search in petabase-scale sequence repositories. Nature pp. 1–9 (2025)

[17] Katz, K., Shutov, O., Lapoint, R., Kimelman, M., Brister, J.R., OSullivan, C.: The sequence read archive: a decade more of explosive growth. Nucleic acids research 50(D1), D387–D390 (2022)

[18] Katz, K.S., Shutov, O., Lapoint, R., Kimelman, M., Brister, J.R., OSullivan, C.: Stat: a fast, scalable, minhash-based k-mer tool to assess sequence read archive next-generation sequence submissions. Genome biology 22(1), 270 (2021)

[19] Lemane, T., Lezzoche, N., Lecubin, J., Pelletier, E., Lescot, M., Chikhi, R., Peterlongo, P.: Indexing and real-time user-friendly queries in terabyte-sized complex genomic datasets with kmindex and ora. Nature Computational Science 4(2), 104–109 (2024)

[20] Marchet, C.: Advances in colored k-mer sets: essentials for the curious (2024), URL https://arxiv.org/abs/2409.05214

[21] Rahman, A., Dufresne, Y., Medvedev, P.: Compression algorithm for colored de bruijn graphs. Algorithms for Molecular Biology 19(1), 20 (2024)

[22] Saitou, N., Nei, M.: The neighbor-joining method: a new method for reconstructing phylogenetic trees. Molecular biology and evolution 4(4), 406–425 (1987)

[23] Uhlmann, J.K.: Satisfying general proximity / similarity queries with metric trees. Information Processing Letters 40(4), 175–179 (1991), ISSN 0020-0190, 10.1016/0020-0190(91)90074-R, URL https://www.sciencedirect.com/science/article/pii/002001909190074R

[24] Yianilos, P.N.: Data structures and algorithms for nearest neighbor search in general metric spaces. In: Proceedings of the Fourth Annual ACM-SIAM Symposium on Discrete Algorithms, p. 311321, SODA ‘93, Society for Industrial and Applied Mathematics, USA (1993), ISBN 0898713137

[25] Zhang, T., Gupta, G., Desai, A., Shrivastava, A.: Identity with locality: An ideal hash for gene sequence search. In: Proceedings of the 31st ACM SIGKDD Conference on Knowledge Discovery and Data Mining V. 1, pp. 1972–1983 (2025)

