## Supplementary Materials for "Lossless compression of k-mer matrices enabling random row access"

### Supplementary Material for "Lossless compression of $k$-mer matrices enabling random row access"

#### S1 Block size effects on compression

Figure S1 provides compressed sizes for various block sizes. We observe that the output sizes of the Human gut metagenome and *S. enterica* data sets behave unexpectedly. We expected the output size to decrease as the block size increased, since the compressor is theoretically more effective with a larger context. We guess that the peaks of the Human gut metagenome curves are due to chance. For the *S. enterica* dataset, we see that $\pi$-compression reaches a local optimum at a block size of 256 KB; further increasing the block size appears to capture greater redundancies.

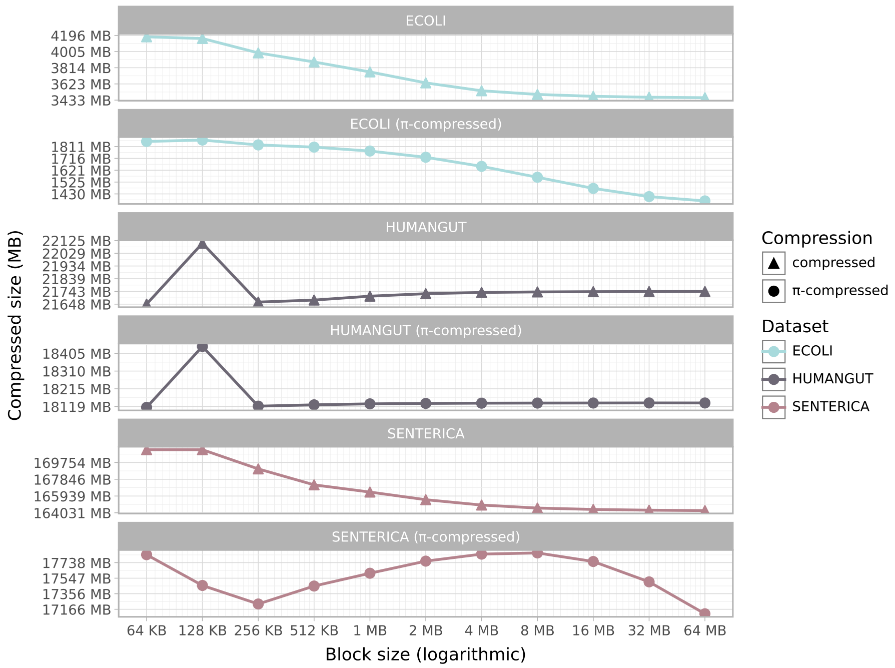


Fig. S1: Resulting index size whether matrices are compressed or $\pi$-compressed according to block size. Zstd's window size was set to the corresponding block size. The subsampling size was set to 10,000.

#### S2 On efficient column operations in row-major bit matrices

In practice, $k$-mer matrices are row-major. This means that column bits must be retrieved individually from each row, which makes column operations inefficient. So, we first apply a bit matrix transposition on the $k$-mer matrix $M$ using an SSE2 approach [2] to make column bits contiguous in memory, giving $M^T$.

**Computing Hamming distance between two columns** Any Hamming distance is computed using a modified AVX2 Harvey-Seal popcount [3], which introduces XOR between two matrix-transposed columns (*i.e.* on rows of $M^T$).

**Practical way to apply a permutation on bit matrix columns** For applying a given permutation on columns, we can either apply such a permutation at the row-level by permuting bits in each row or at the matrix-level by moving columns. But $k$-mer matrices can have up to 100k columns or more. At the row level, there is no efficient way to permute such bit sequences. It would imply permuting each bit individually in each row, which yields roughly 80 MB/s (as shown in Figure S3). Given the input problem size, we cannot apply solutions such as AVX-512 dynamic bit shuffle [1] or bit permutation methods described in [4], which are meant for much smaller instances, limited to a few hundred bits. This explains why, as depicted in Figure 1, we first transpose the bit matrix. Afterward, permutation $\pi$ is applied in $\mathcal{O}(N)$ by solving its disjoint cycles and moving transposed columns. Finally, the matrix is transposed back. Formalization is defined as follows.

**Definition S1.** *(permuting columns with double transposition) Let $\mathcal{M}$ be a bit matrix. Let $\pi_c(\mathcal{M})$ and $\pi_r(\mathcal{M})$ be two functions that apply a permutation $\pi$ respectively to columns and rows of $\mathcal{M}$.*

Note that it is possible to compute $\pi_c(M)$ by using only $\pi_r$ and a matrix transposition ($^T$) as $\pi_c(M) = (\pi_r(M^T))^T$.

**Efficient column permutation with double block transposition** Once the permutation $\pi$ is determined for the columns of a matrix $M$, we want to compute $\pi_c(M)$. In practice (see Figure 1 in the main text), we apply these three steps:

1. Transpose $M$ to $M^T$.
2. Apply an in-place permutation of rows (transposed columns) of $M^T$, obtaining $\pi_r(M^T)$.
3. Transpose the obtained matrix $\pi_r(M^T)$, obtaining $(\pi_r(M^T))^T$, thus obtaining $\pi_c(M)$.

However, bit matrix transposition is not memory-efficient, as it involves numerous random memory accesses that can lead to substantial cache misses. Yet any matrix transposition can be expressed as a block-matrix transposition. We split the input matrix $M$ horizontally into several sub-matrices $M_0, M_1, \ldots, M_X$. We apply the previously presented column reordering procedure to each submatrix individually (see Figure 1). For each $M_i$, one computes $\pi_c(M_i)$ by computing $(\pi_r(M_i^T))^T$. We call this a reordering with "double block transposition".

**Bitpacking vs. double block transposition** As depicted in Figure S2, double block transposition is faster than bitpacking by 2-5× factor depending on matrix dimensions, enabling tractable throughput as shown in Figure S3. Even though bitmatrix block transpositions are only data preprocessing and postprocessing steps that do not perform any reordering, they enable SIMD vectorization of column reordering and are more CPU cache-friendly. Note that a throughput of less than 100 MB/s would be a major bottleneck, making the whole pipeline inefficient.

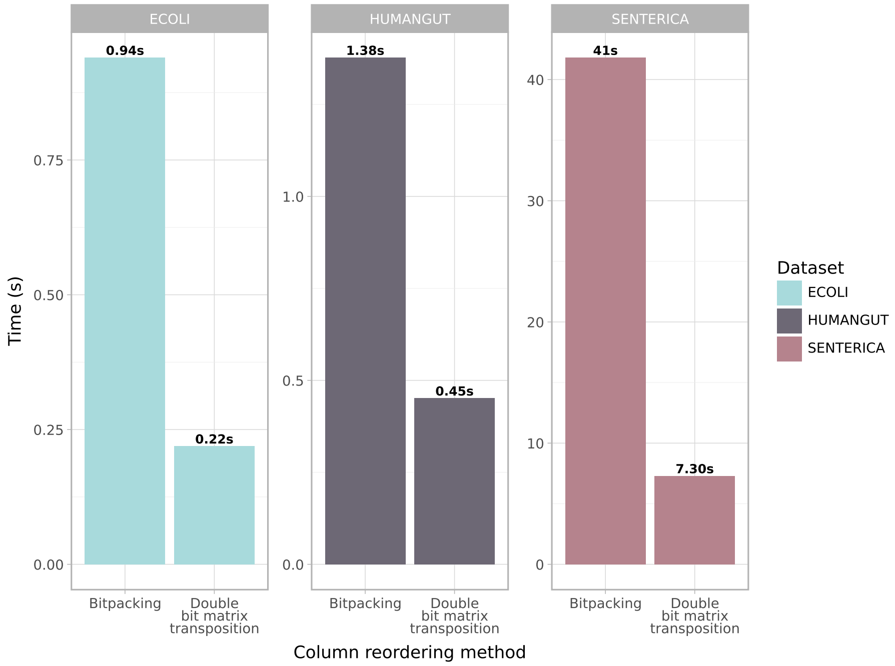


Fig. S2: Bit matrix column reordering time according to method and matrix type. Transposed block size was set to 8MB.

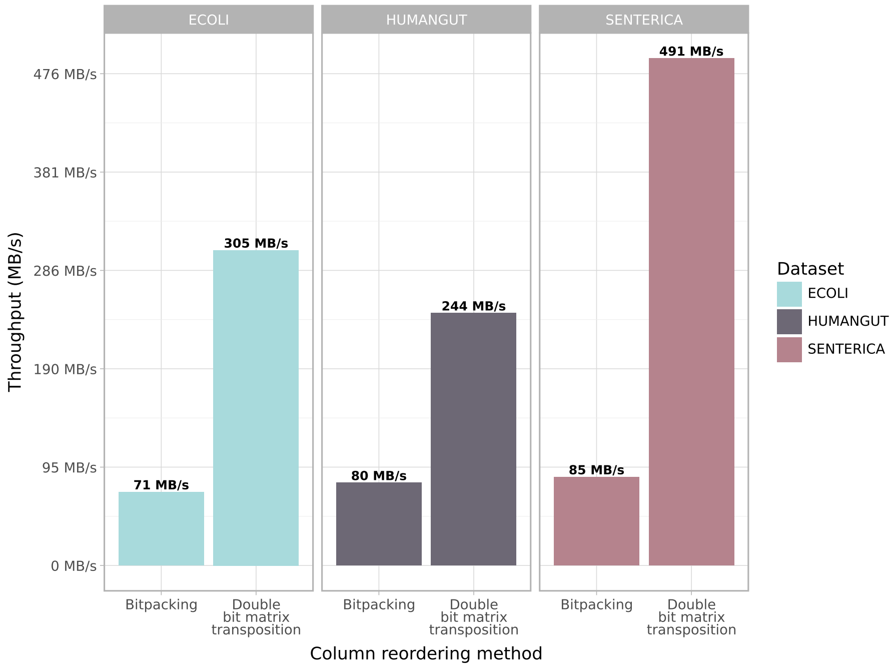


Fig. S3: Column reordering method mean throughput. Transposed block size was set to 8 MB.

#### S3 Computing the Hamming distance between columns

We use the Hamming distance for estimating the similarity between two columns. This distance has several advantages. First, the Hamming distance validates the triangle inequality property needed by `vp-tree`. Second, once this distance is rescaled to the $[0;1]$ range (see Definition S2), restricting attention to row bits, it gives the probability that a bit from column A does not share the same value as the corresponding bit from column B, in other words, the probability of breaking a bit-run within a bit vector. Finally, a Hamming distance is fast to compute, as it can be computed using a XOR and a popcount. Nonetheless, our $k$-mer matrices are row-major, so any operation on columns is slow. We give more details about how we deal with such constraints in Section S2.

**Definition S2 (Scaled Hamming distance).** *Let $|A|$ be the length of bit-string $A$. The scaled Hamming distance is* $d^s_H(A,B) := \frac{d_H(A,B)}{|A|} = \frac{H(A \oplus B)}{|A|}$.

#### S4 Row subsampling for faster distances computations

In a $k$-mer matrix, each column can reach several million bits. Hence, the computation time for Hamming distance on such bit sequences can be unnecessarily long. We solve this issue by subsampling the bits used for the Hamming distance computations. We assume that $k$-mers are randomly and uniformly distributed in $M$ rows. In the case of `kmindex`, $k$-mers were hashed using `xxHash`[1]. This hash function has well-defined dispersion, and its bits can be considered independent. Hence, taking the first $h$ rows as a subsample, generating the matrix $M_h \in \{0,1\}^{N \times h}$ is an unbiased sequence of column sketches.

#### S5 The `vp-tree` space and time complexities

Given $N$ elements, a vantage-point tree can be constructed in time $\mathcal{O}(N \log N)$. It provides an average $\mathcal{O}(\log N)$ time complexity for determining the nearest neighbor of an element, while in the worst case, this operation takes $\mathcal{O}(N)$ time. Hence, in the worst case, the full Nearest Neighbor algorithm has $\mathcal{O}(N^2)$ time complexity (just as a naive implementation), while in the best case it has $\mathcal{O}(N \log N)$ time complexity.

[1] `https://github.com/Cyan4973/xxHash`

**Algorithm S1** `masked vp-tree`: nearest unvisited neighbor search

**Require:**
$N \in \mathbb{N}^*$ ▷ Number of vertices
$\ell \in [\![0; N-1]\!]$ ▷ Vertex identifier
$\alpha \in [\![0; N-1]\!]$ ▷ Nearest unvisited neighbor identifier
$\tau \leftarrow +\infty$ ▷ Nearest unvisited neighbor distance
alreadyAdded $\in \{\bot, \top\}^N$

```
function NearestUnvisitedNeighbor(node, ℓ, α, τ)
    if node = null then
        return
    end if

    L ← node.left
    R ← node.right
    μ ← node.threshold
    p ← node.pivot
    d ← d_H(ℓ, p)                                   ▷ Hamming distance between ℓ and p

    if (¬alreadyAdded[p]) ∧ (d < τ) then
        τ ← d
        α ← p
    end if

    if d < μ then                                   ▷ Branch-and-bound
        if (L ≠ null) ∧ (¬L.masked) ∧ (d − τ ≤ μ) then
            NearestUnvisitedNeighbor(L, ℓ, α, τ)
        end if
        if (R ≠ null) ∧ (¬R.masked) ∧ (d + τ ≥ μ) then
            NearestUnvisitedNeighbor(R, ℓ, α, τ)
        end if
    else
        if (R ≠ null) ∧ (¬R.masked) ∧ (d + τ ≥ μ) then
            NearestUnvisitedNeighbor(R, ℓ, α, τ)
        end if
        if (L ≠ null) ∧ (¬L.masked) ∧ (d − τ ≤ μ) then
            NearestUnvisitedNeighbor(L, ℓ, α, τ)
        end if
    end if
end function
```

**Algorithm S2** `masked vp-tree`: masked property update

```
Require:
  N ∈ ℕ*                                          ▷ Number of vertices
  alreadyAdded ∈ {⊥, ⊤}^N

  function UpdateMaskedProperty(node)
      while node ≠ null do
          L ← node.left
          R ← node.right
          p ← node.pivot

          node.masked ← (L = null ∨ L.masked) ∧
                        (R = null ∨ R.masked) ∧
                        alreadyAdded[p]
          if ¬node.masked then
              return
          end if

          node ← node.parent
      end while
  end function
```

#### S6 Implementation

All implementations are available as GitHub repositories. The block compression scheme is available at `https://github.com/AlixRegnier/BlockCompressor`, the library for reordering and compressing the matrices called `kmcomp` is available at `https://github.com/AlixRegnier/kmcomp`, and the libraries were added to `kmindex`, which is available at `https://github.com/tlemane/kmindex/tree/next_realease`. Scripts used to generate results can be found at `https://github.com/AlixRegnier/kmcomp_experiments`.

#### S7 Hardware specification

Measures on experimental datasets were done with a single thread on an Intel(R) Xeon(R) Gold 6338N at 2.20 GHz. We are using 16x 3.84 TB HPE VK003840GZXRH SSDs in RAID 6. We use 4 KB blocks for read measurements and 1 MB blocks for write measurements. We report an average bandwidth of 6055 MiB/s and a latency of 0.33ms for read measures. We report an average bandwidth of 2886 MiB/s and a latency of 177.19ms for write measures.

#### S8 Datasets

Experimental datasets were indexed using `kmindex`, with default values. Results about the gain obtained in the Hamming distances between consecutive columns with and without applying the $\pi$ permutation and about the $\pi_{\text{gain}}$ values are provided in Table S1.

Table S1: Experimental datasets properties. Distance used is the scaled Hamming distance, defined between 0 and 1. Subsampling size was set to 10,000.

| Dataset | Initial mean consecutive distance | New mean consecutive distance | $\pi_{\text{gain}}$ |
|---|---|---|---|
| *E. coli* | 0.06834 | 0.01434 | 4.764 |
| Human gut metagenome | 0.34058 | 0.19990 | 1.704 |
| *S. enterica* | 0.05073 | 0.00173 | 29.257 |

#### S9 Saved bits per second formulas

Following equations give how we calculated compression efficiency when applying compression or $\pi$-compression:

Compression:

$$\frac{\text{raw size} - \text{compressed size}}{\text{compression time}} \tag{S1}$$

$\pi$-compression:

$$\frac{\text{raw size} - \text{compressed size}}{\pi\text{computation time} + \text{compression time (of the permuted matrix)}} \tag{S2}$$

#### S10 Per-stage runtime on experimental datasets

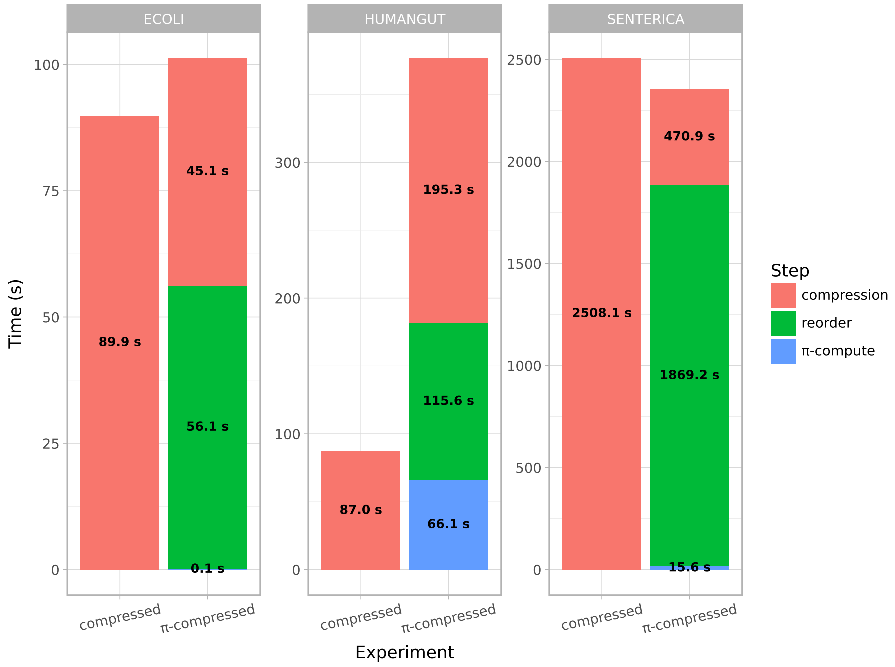


Fig. S4: Stage time whether matrices are compressed or $\pi$-compressed. Subsampling size was set to 10,000. Block size was set to 64 KB.

In Figure S4, we can see that compression times vastly differ when matrices are reordered or not. It can be explained by the fact that, in our current pipeline, we directly compress blocks in RAM as soon as they have been reordered. When $\pi$-compressed, data loading is done during the reordering step. We also guess that disk write speed is important because the better the data is compressed, the faster it is written to disk. Finally, we note that altering data can cause Zstd to select a different compression strategy, in which less complex data may be compressed more quickly. We also report that the computation time of $\pi$ only takes a small part of the total pipeline's cumulative time, except for Human gut metagenome.
Note that compression and reordering steps can be heavily multithreaded.

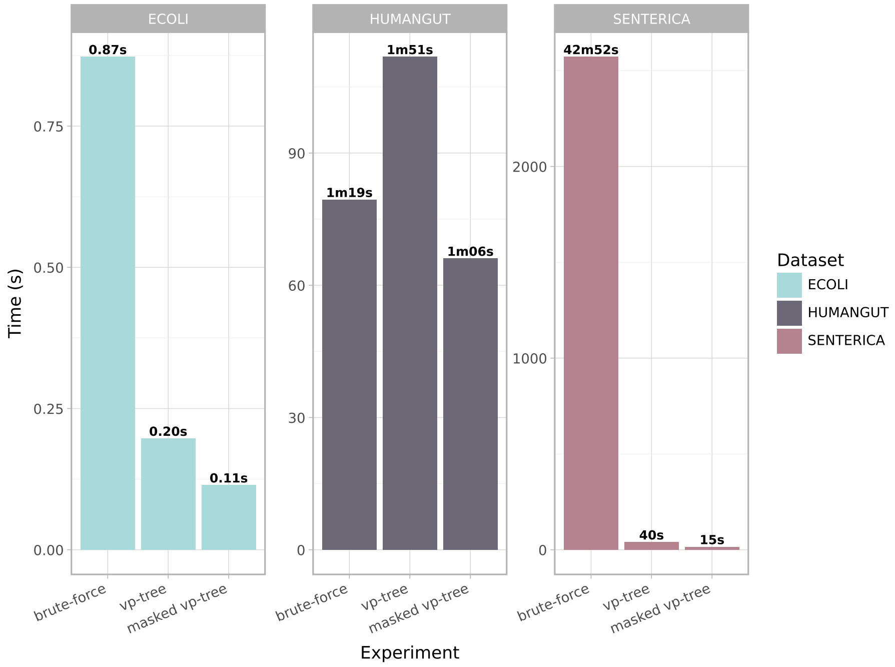


Fig. S5: Permutation ($\pi$) computation time according to method and matrix type. Subsampling size was set to 10,000. Repeated 10 times with a mean relative standard deviation of 9.5%.

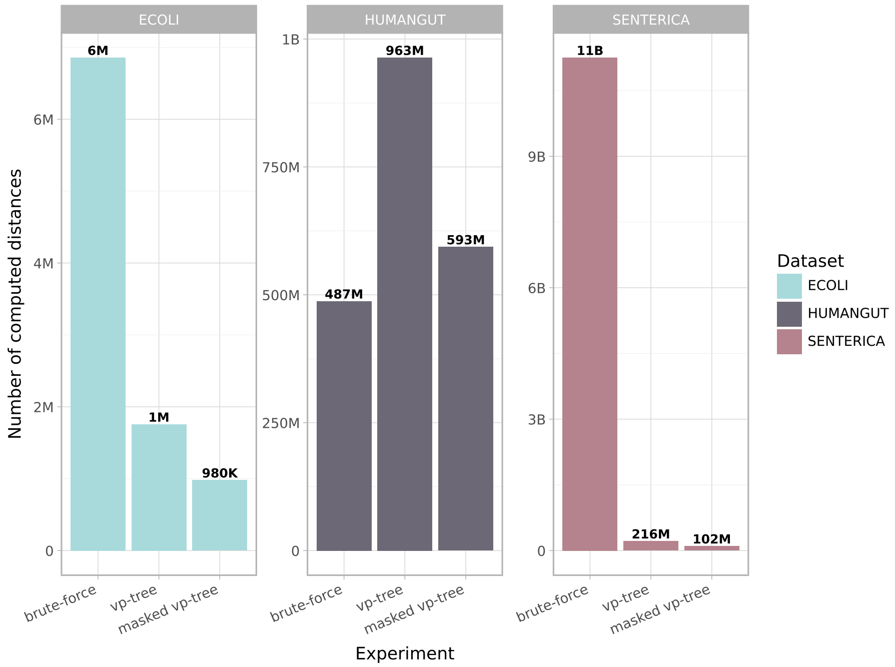


Fig. S6: Number of computed distances according to method and matrix type.

Figures S5 and S6 respectively report the runtime and number of computed distances for $\pi$ under three strategies: brute-force, `vp-tree`, and `masked vp-tree`. The two figures jointly illustrate the strong correlation between distance computation count and overall runtime.

#### S11 Varying subsampling size

The more subsampled rows, the better the distance accuracy. Note that the permutation computation time mainly depends on the number of samples, the number of computed distances, and the subsampling size. According to Figure S7, it seems that subsampling size has an intrinsic effect on the number of computed distances depending on the dataset. Indeed, as shown in Figure S8, the number of computed distances is plotted against subsampling size. For the *E. coli* dataset, the number of computed distances increases logarithmically when distance accuracy increases linearly. For the Human gut metagenome, the number of computed distances rapidly reaches the number of pairwise distances. For the *S. enterica* dataset, increasing the subsampling size appears to improve `masked vp-tree` search space pruning during nearest-neighbor queries. Figure S9 shows that the larger the subsampling size, the smaller the compressed size. However, the compressed size appears to decrease logarithmically, meaning that increasing the subsampling size will increase $\pi$ computation time linearly, while the compression will only be slightly improved. Thus, the subsampling size parameter is a trade-off between time and output compression. According to our experiments, a subsampling size between 1,000 and 20,000 seems to be a reasonable parameter range if one needs either fast reordering or better compression.

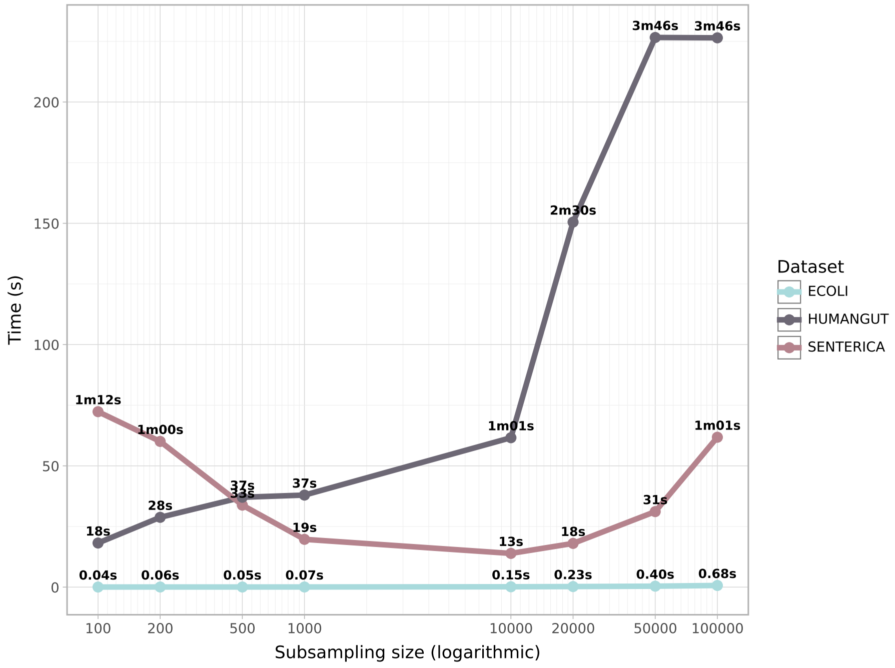


Fig. S7: Permutation ($\pi$) computation time according to subsampling size and matrix type.

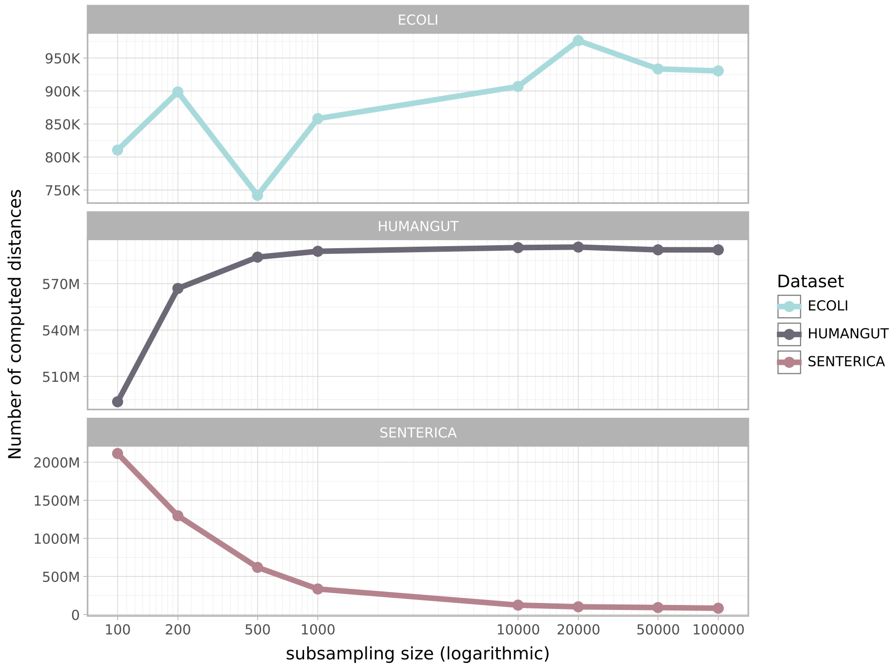


Fig. S8: Number of computed distances according to subsampling size and matrix type.

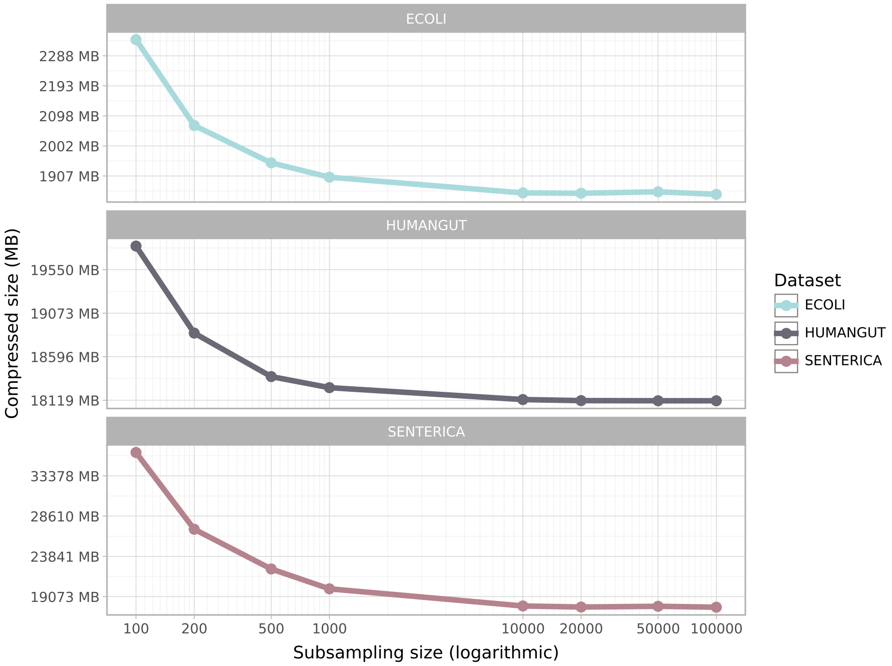


Fig. S9: Resulting $\pi$-compressed indexes size according to subsampling size. Block size was set to 64 KB.

#### S12 Query time according to block size

For queries, Figure S10 shows that smaller blocks yield faster query times. This is expected, as smaller blocks reduce the likelihood of decompressing data irrelevant to the query. We anticipated that the overhead of switching between blocks would be a limiting factor; however, this does not appear to be the case. We attribute this to Zstd's ability to reuse decompression data structures across blocks, thereby avoiding repeated memory allocation. Overall, these results suggest a trade-off between compression ratio, block size, and query size. According to our results, more investigation is needed to understand what would happen on smaller blocks.

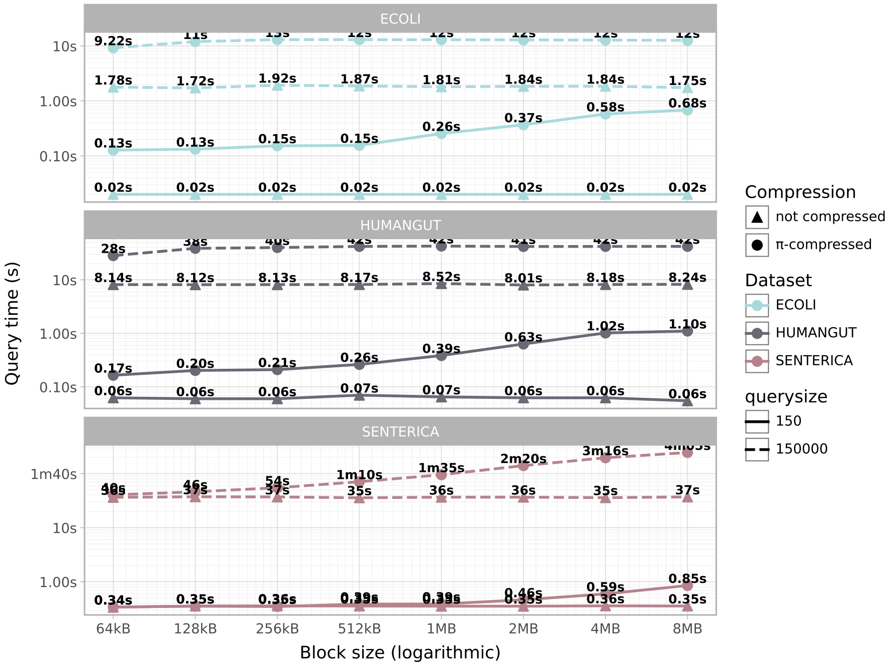


Fig. S10: Mean query time whether matrices are not compressed or $\pi$-compressed according to block size. Zstd's window size was set to the corresponding block size. Repeated 8 times with a mean relative standard deviation of 2.67%.

#### S13 Testing $\pi_{gain}$ on Logan-search $k$-mer matrices

Figure S11 shows results comparing $\pi_{gain}$ estimation compared to actual compression ratios. We used $k$-mer matrices with at least 4096 columns to compute the $\pi_{gain}$ formula, as this metric is meaningful only when there are sufficient columns. The linear regression equation is $0.414 \times x + 0.596$ with a $R^2$ of $\approx$ 0.94. In Figure S12, we show that in a given $k$-mer matrix for bacteria from `Logan-Search` index, only 3.2% of the accessions were correctly classified as bacteria.

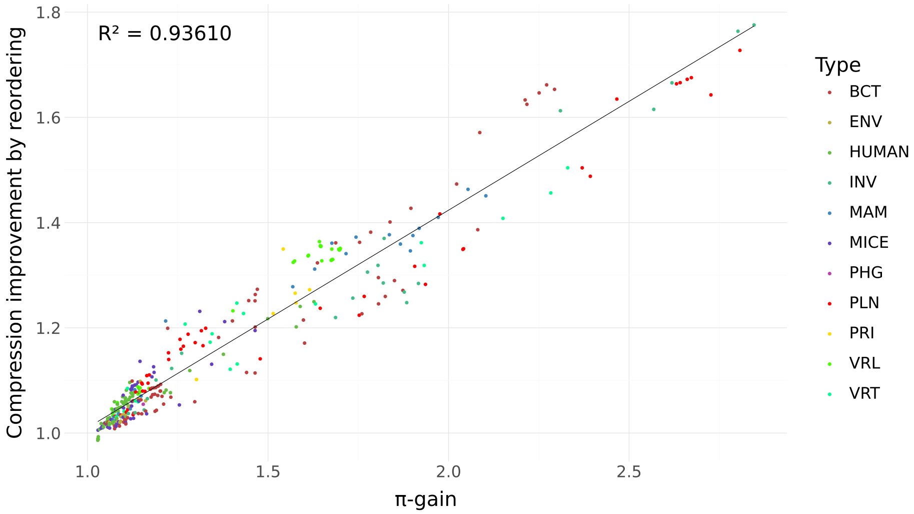


Fig. S11: Linear regression of $\pi_{\text{gain}}$ predicting $\pi$-compression improvement over compression. 349 $k$-mer matrices from `Logan-Search` containing at least 4096 columns were used to compute the linear regression. The colors correspond to the types of genomic data indexed in the matrices. The type of data represented is the one used in `Logan-Search`, whose correspondences are given in Table S2

Table S2: Abbreviations used in Logan-Search sub-indexes name

| Abbreviation | Taxon |
|---|---|
| BCT | Bacteria |
| ENV | Environmental sample |
| HUMAN | Human |
| INV | Invertebrate |
| MAM | Mammal |
| MICE | Mice |
| PHG | Phage |
| PLN | Plant |
| PRI | Primate |
| ROD | Rodent |
| UNKNOWN | Unknown |
| VRL | Viral |
| VRT | Vertebrate |

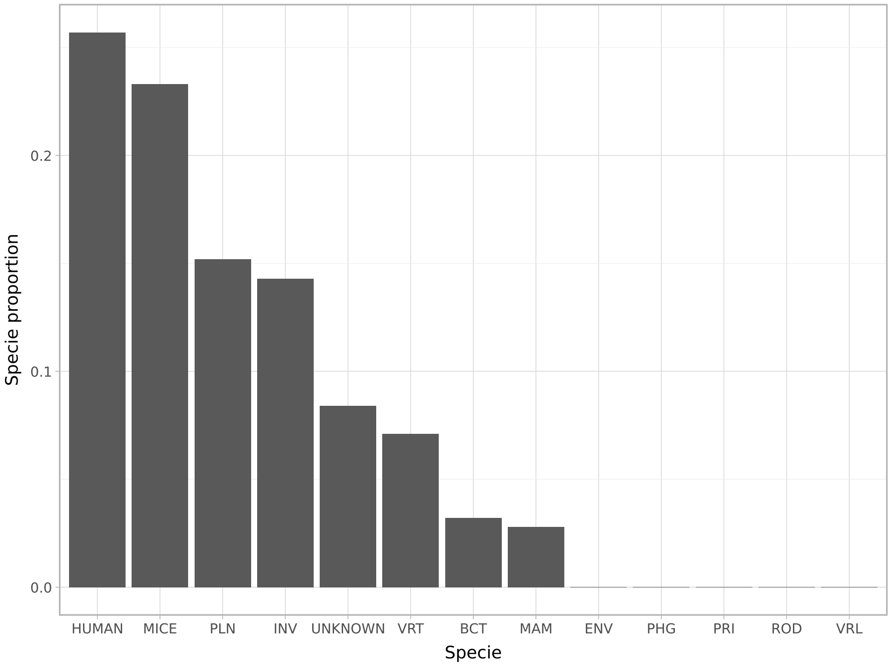


Fig. S12: NCBI annotated classification of 131072 accessions in `Logan-Search` that are annotated as `BCT`. Accession taxonomic divisions have been retrieved using APIs from NCBI SRA and ENA taxonomy.

### Supplementary Material references


[1] Lemire, D.: Dynamic bit shuffle using AVX-512 (2023), URL https://lemire.me/blog/2023/06/29/dynamic-bit-shuffle-using-avx-512/

[2] Misha Sandberg: What is SSE !@# good for? Transposing a bit matrix (2011), URL https://mischasan.wordpress.com/2011/07/24/what-is-sse-good-for-transposing-a-bit-matrix/

[3] Muła, W., Kurz, N., Lemire, D.: Faster population counts using avx2 instructions. The Computer Journal **61**(1), 111–120 (2018)

[4] Sirrida: Bit permutations (2020), URL http://programming.sirrida.de/bit_perm.html
